# Anatomical diversification of a skeletal novelty in bat feet

**DOI:** 10.1101/490854

**Authors:** Kathryn E. Stanchak, Jessica H. Arbour, Sharlene E. Santana

## Abstract

Neomorphic, membrane-associated skeletal rods are found in disparate vertebrate lineages, but their evolution is poorly understood. Here we show that one of these elements—the calcar of bats (Chiroptera)—is a skeletal novelty that has anatomically diversified. Our comparisons of evolutionary models of calcar length and corresponding disparity-through-time analyses indicate that the calcar diversified early in the evolutionary history of Chiroptera, as bats systematically radiated after evolving the capacity for flight. We find interspecific variation in a variety of anatomical parameters of probable importance for calcar function, which suggests that adaptive advantages provided by the calcar led to its anatomical diversification. In addition to overall length, we find that the calcar varies among bats in its tissue composition, and a synovial joint is present at the articulation between the calcar and the calcaneus ankle bone in some species. This suggests the calcar has a kinematic functional role. Our results demonstrate that novel skeletal additions can become integrated into vertebrate body plans and subsequently evolve into a variety of forms, potentially impacting clade diversification by expanding the available morphological space into which organisms can evolve.

## INTRODUCTION

Enigmatic, neomorphic anatomical elements are scattered throughout the paleontological and neontological records of vertebrate evolution (Hall 2015). Recent fossil discoveries have raised interest in one specific type of novel skeletal structure: the “styliform” elements of vertebrates that use membranes to glide or fly (Fig. 1). This group of skeletal elements comprises the calcar of bats (Schutt and Simmons 1998), the styliform cartilages of gliding rodents and one marsupial (Coster et al. 2015, Kawashima et al. 2017, Johnson-Murray 1987, Jackson 2012), the pteroid of pterosaurs (Bennett 2007), and was recently expanded to include the styliform element of *Yi qi*, a maniraptoran theropod dinosaur (Xu et al. 2015) and the calcar of *Maiopatagium furculiferum*, a haramiyid mammaliaform (Meng et al. 2017). Since these skeletal rods are now known from disparate tetrapod lineages, they seem less like evolutionary oddities than consequential skeletal novelties characteristic of membranous body plans. The literature on most of these structures is limited to osteological descriptions, so much is still unknown about their function, origin, and diversification. The pterosaur pteroid has been the focus of several studies and has generated debates on its anatomy and function (summarized in Witton 2013), but although the Pterosauria comprises a taxonomically diverse clade in which to explore pteroid variation, the lack of extant successors in the lineage restricts detailed anatomical and functional studies. In contrast, another of these neomorphic styliform elements—the bat calcar—is widespread across extant bats, making it an ideal model system for gaining a better understanding of the evolution of membrane-bound skeletal rods, and more generally, the evolution of neomorphic skeletal elements.

**Figure 1.**
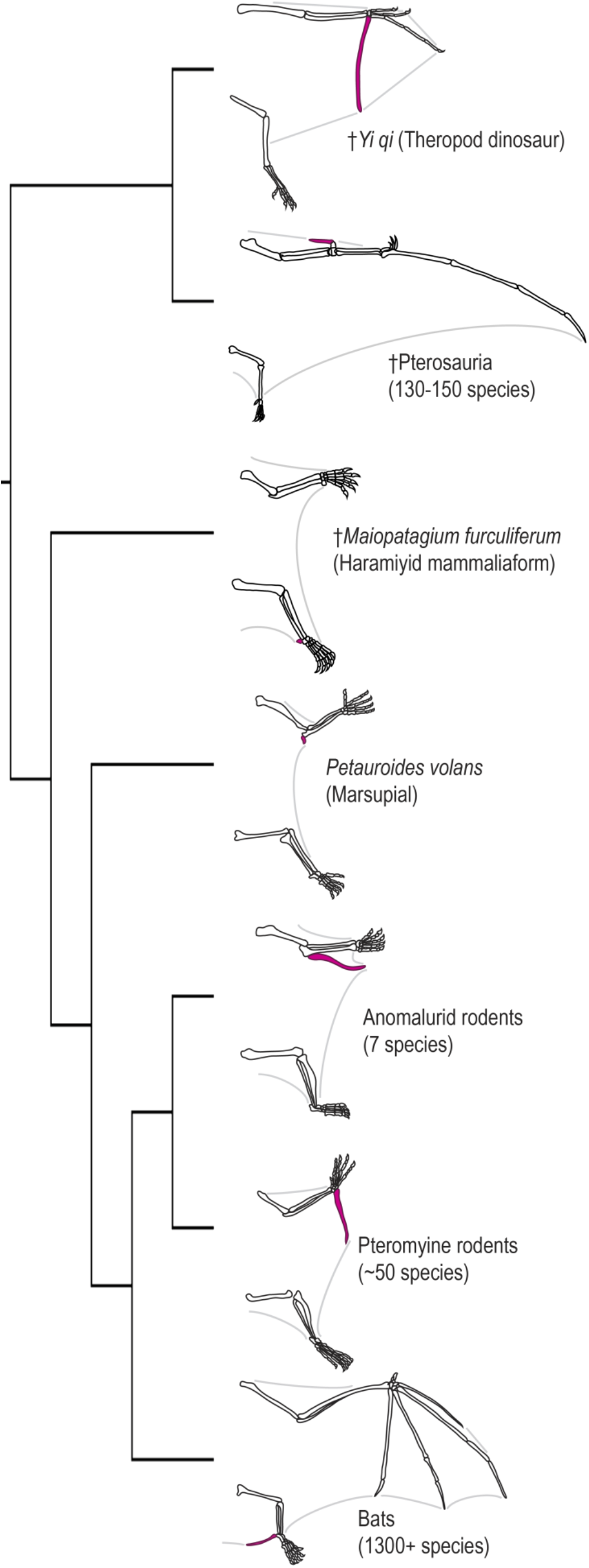
Neomorphic skeletal rods have evolved multiple times in vertebrates with gliding or flying membranes. These structures are indicated in pink in the schematic drawing. Drawings based on Xu et al. 2015, Meng et al. 2017, Bennett 2007, Witton 2013, Coster et al. 2015, Kawashima et al. 2017, Johnson-Murray 1987, Jackson 2012.

Bats (Chiroptera) are systematically, morphologically, and ecologically diverse (Simmons 2005, Fenton and Simmons 2015, Kunz and Fenton 2005). The calcar articulates with the calcaneus bone in the bat ankle and extends into the membrane that spans between the two hindlimbs (Vaughan 1970; Fig. 1). The calcar abruptly appears in the early bat fossil record (*Onychonycteris finneyi*, Onychonycteridae, Green River Formation, WY, USA; ~52.5 Ma; Simmons et al. 2008) and, based on its ubiquity among extant bats, seems to have become fixed as part of the bat wing structure. It is typically described as a bony or cartilaginous element, although histological studies to date have confirmed only the presence of cartilaginous tissue with varying levels of mineralization (Schutt and Simmons 1998, Adams and Thibault 1999, Czech et al. 2008, Stanchak and Santana 2018). Because bats are morphologically diverse and cartilage can be a precursor of bone, it has been hypothesized that the calcars of some bat species might be composed of bony tissue (Adams and Thibault 1999).

The calcars of the Old World fruit bats (Pteropodidae) are known to be different from those of the other bats. Pteropodid calcars are described as inserting on the tendon of the gastrocnemius muscle rather than articulating with the calcaneus and are consequently hypothesized not to be homologous to the calcars of other bats (Schutt and Simmons 1998, Kobayashi 2017). In previous phylogenetic hypotheses, Pteropodidae was considered the sister clade to all of the other bat families, which were collectively referred to as the “microchiroptera.” However, after the phylogeny of Chiroptera was revised using molecular data, non-pteropodid bats were rendered paraphyletic (Teeling et al. 2005). As a consequence, the hypothesis of a lack of homology between the pteropodid calcar and that of the “microbats” became a less-parsimonious explanation than that of a homologous calcar across Chiroptera.

In all animal clades with styliform elements, including bats, the evolution of membrane-bound limbs and a new locomotor mode (flight or gliding) allowed entry into new ecological space: the aerosphere. The bat fossil record demonstrates early taxonomic diversification coupled with a rapid expansion of their geographic distribution (Smith et al. 2012). The earliest known bats, onychonycterids, have been found on both the North American and Eurasian Eocene land masses (Hand et al. 2015). By the end of the Eocene, bats are known from six continental landmasses (Smith et al. 2012, Hand et al. 2015). *Onychonycteris*.*finneyi*, which possessed the earliest known calcar, also had the most transitional bat postcranial skeleton found to-date, with limb proportions between those of bats and non-volant mammals (Simmons et al. 2008). Based on its presence in the oldest bat fossils, the calcar may be part of the suite of adaptations that allowed bats to functionally and ecologically radiate into varied niches after their initial invasion of the aerosphere. If so, we predict that (1) bat calcars will be morphologically diverse in trait parameters that theoretically affect function, and (2) calcar morphological diversification will reflect the rapid early diversification of Chiroptera, as suggested by the fossil record.

In this paper, we assess and describe the anatomical diversification of the calcar across the radiation of bats to test the predictions outlined above. We integrate a variety of methods to analyze calcar anatomy across a broad sample of bat species spanning diverse ecologies. First, we examine the variation in length of the calcar across Chiroptera and test different models of calcar evolution to reveal the macroevolutionary patterns and potential underlying processes that characterize calcar diversification. Then, we more closely investigate the anatomical diversity of the calcar with micro-Computed Tomography (μCT) scans to assess its status as a novel skeletal addition rather than another type of skeletal modification (e.g., a repeated tarsal element), and we integrate data from both μCT scans and histological sections to test the hypothesis that the calcar has histologically diversified. Finally, we combine gross dissections and diffusible iodine-based contrast-enhanced μCT (diceCT; Gignac et al. 2016) for the visualization of soft tissue to re-evaluate the hypothesis that the pteropodid calcar is not homologous to the calcar of other bats. Collectively, these studies rigorously assess the scope and scale of bat calcar evolution.

## MATERIAL AND METHODS

### Calcar Length Measurements and Macroevolutionary Analyses

The length of a rod or shaft is one parameter that determines its ability to resist bending under an applied load (Hibbeler 2007). Bat calcars generally take a rod-shaped form, so comparisons of calcar length are informative about the potential functional importance of the calcar across bats. A single observer (KES) made caliper measurements of calcar, tibia, and forearm (i.e., radius) lengths of 1-9 fluid-preserved specimens representing 226 species and all recognized families within Chiroptera. In total, the sample included 1,396 specimens with an average of 6 specimens per species. A list of museum specimens is provided as a spreadsheet in the Supporting Information. By measuring intact, fluid-preserved specimens, we ensured that any thin, cartilaginous portions of the calcar were present and measured. We rounded caliper measurements to the nearest 1mm to reflect imprecision in measuring skeletal features from external examination of intact specimens. Because we based all measurements on external examination of specimens, it is possible that a very small, not externally evident calcar resulted in assigning a value of zero calcar length to some individuals (e.g., see Results regarding *Rhinopoma hardwickii*). We did not include fossil bat species in our sample because few postcrania are present in the bat fossil record and some extant calcars are unmineralized, so we would not be able to confirm the absence of a calcar for any bat fossil species.

For each specimen, we calculated the ratio of the calcar length divided by either the tibia or the forearm length and then averaged these ratios across all specimens for a particular species to derive a unitless measure of hindlimb skeletal proportions to compare across species. We visualized the calcar-to-tibia length ratio character states on a pruned version of a relatively recent chiropteran phylogeny (Shi and Rabosky 2015) using the “fastAnc” method of the “contMap” function (Felsenstein 1985, Revell 2013) from the *phytools* v.0.6 package (Revell 2012) in R v.3.4.3 (R Core Team 2017; all analyses were performed in the same version of R). We also calculated the residuals of phylogenetic generalized least squares regressions (pgls) of mean calcar length on mean tibia or mean forearm length assuming a Brownian motion correlation structure using the “phyl.resid” function (Revell 2009, Revell 2010) from the *phytools* v.0.6 R package (Revell 2012). While the calcar-to-tibia length ratio is more intuitively relevant to calcar biomechanics and function-even beyond its use for size normalization-we used both the tibia and forearm ratios and pgls residuals in subsequent evolutionary analyses so that we could better interpret the effect of variable transformations on our model fits. In addition, we repeated all of the following analyses for datasets that did not include the species for which we measured zero calcar length, and from which we excluded the Pteropodidae due to their differing calcar anatomy. All data used in analyses are provided as a spreadsheet in the Supporting Information.

To gain insight on the evolutionary processes that may have led to extant calcar diversity, we fit three models of evolution (Brownian motion, early burst, and single-peak Ornstein-Uhlenbeck) to the calcar length ratios and pgls residuals using the “fitContinuous” function in the *geiger* v.2.0.6 R package (Harmon et al. 2007, Pennell et al. 2014). Brownian motion (BM) models a “random-walk” process in which the variance of a trait increases linearly through time (as defined in evolutionary modeling by the evolutionary rate parameter σ^2^). It is often used to test the hypothesis of trait evolution under a drift or other random process (Felsenstein 1973). The early burst (EB) model is used to test a niche-filling hypothesis consistent with an adaptive radiation; the rate at which a trait diversifies decreases with declining ecological opportunity after an initial, rapid “early burst” of diversification (Blomberg et al. 2003, Harmon et al. 2010). The EB model is parameterized by the initial evolutionary rate (σ^2^) and a parameter for the exponential change in evolutionary rates through time (a), such that when a = 0 the EB model reduces to the BM model and when a < 0 evolutionary rates decrease as time progresses. An Ornstein-Uhlenbeck (OU) process is used to model an evolutionary process in which some restoring force (e.g., selection; parameterized by α) restrains a trait value (θ) through time (Hansen 1997, Butler and King 2004). As implemented here, the model assumes a single optimal trait value that is equal to the root ancestral state of the trait (parameterized by z_0_ in all models). We compared these three models using small sample size-corrected Akaike weights (*w*_AICc_). If the calcar underwent an early morphological diversification as the first bats systematically radiated, we expected to find the highest support for the EB model.

To visualize and quantify the tempo of calcar length evolution, we performed a disparity-through-time analysis using the “dtt” function (Harmon et al. 2003, Slater et al. 2010) from the *geiger* v.2.0.6 R package (Harmon et al. 2007, Pennell et al. 2014) to calculate the mean morphological disparity of each subtree in the pruned phylogeny using the average squared Euclidean distance among all pairs of points. We plotted this curve against a null distribution created by using the same procedure on a set of 1,000 simulations across the pruned phylogeny assuming a BM model of evolution of the relative calcar lengths. We used the morphological disparity index (MDI) to quantitatively compare subclade disparity in relative calcar length with the disparity expected under a BM model (Harmon et al. 2003, Slater et al. 2010). We determined the significance of the MDI by the frequency at which a calculated MDI between the data set and each simulation trial was greater than zero. A negative MDI value indicates that disparity is partitioned more strongly among early divergence events, with more recent subclades each representing only a small portion of the total morphological diversity of the clade than expected under a constant-rate, random walk process (e.g., BM; Harmon et al. 2003, Slater et al. 2010). Positive MDI values may be indicative of selective constraint or increasing evolutionary rates, where each recent subclade is more likely to represent a greater proportion of trait space (López-Fernández et al. 2013). A negative MDI supports a hypothesis of early, rapid morphological diversification prior to a period of relative stasis until the present day (Slater et al. 2010). To more rigorously assess the prediction of early disparification, we also performed disparity-through-time analyses in which we compared the results from our data against simulated results generated under an EB model of evolution (Slater and Pennell 2014).

### CT Scanning

To examine calcar anatomy in the context of other ankle and foot bones across bat species, we dissected and μCT scanned one foot of each of 19 fluid-preserved bat specimens representing 13 families within Chiroptera. We also μCT scanned three whole (non-dissected) fluid-preserved specimens representing three additional bat families (Appendix S1) for a total sample of 22 species representing 16 families. These specimens were sourced from museum collections, research collections in the Santana Lab and the Herring Lab at the University of Washington, and the Lubee Bat Conservancy. We segmented (digitally-dissected) the tarsals, the calcar, and other accessory ossicles in each μCT scan using Mimics v.19 (Materialise). The resulting 3D renderings allowed us to compare tarsal osteology across our samples in unprecedented detail.

Previous studies of pteropodid calcar anatomy describe a calcar that inserts on the tendon of the gastrocnemius muscle. This tendon then inserts on the calcaneal tuberosity. In contrast, calcars of the paraphyletic “microbats” articulate directly with the calcaneus. To better assess the soft tissue morphology of the calcars in Pteropodidae, we used diffusible-iodine contrast-enhanced μCT scanning (diceCT, Gignac et al. 2016) and conventional μCT scanning to image the feet of the three pteropodid species in our sample. For diceCT scanning, we placed each fluid-preserved specimen in a solution of Lugol’s iodine (3% total solute) for two to three days prior to CT scanning. The iodine solution increases the x-ray opacity of soft tissue—particularly muscle—in the sample, allowing for the visualization of this tissue in the μCT scan. Then, we dissected each of the pteropodid feet to further assess the connection between the calcar spur and the calcaneus ankle bone. A list of scanned specimens and μCT scanner settings is provided in Appendix S1.

### Histology

We used both the μCT scans and histological sections of the dissected specimens to compare calcar tissue composition across 18 bat species (Appendix S1). Calcified calcar samples were first decalcified in 14% EDTA aqueous solution neutralized with ammonium hydroxide. Because we had difficultly completely decalcifying some samples in EDTA, we transferred them to 5% aqueous formic acid for further decalcification. We then dehydrated, cleared, and embedded all samples in paraffin wax. We sectioned each paraffin block at 5-8 micrometers with a Leica RM2145 microtome, mounted the sections to slides, then cleared, rehydrated, and stained the sections using either modified Mayer’s hematoxylin and Mallory’s triple connective tissue stain (Humason 1962) or Weigert’s iron hematoxylin and fast green/safranin O. For all samples, we determined calcar tissue composition by cell and substrate morphology, not by stain color. We imaged the sections with a Nikon Eclipse E600FN compound microscope and an AmScope MU300 camera.

## RESULTS

The calcar exhibits extensive anatomical diversity across Chiroptera. Calcars range from not externally visible (a length of zero) to considerably longer than the tibia (Fig. 2). We found strong support (*w*_AICc_ > 0.99) for the EB model of morphological evolution for calcar length relative to tibia length in all model comparisons that included pteropodid bats in the sample (Table 1). All OU models collapsed to BM models, so only model results for BM and EB models are shown. Support for the EB model decreased for the sample that did not include Pteropodidae. Disparity-through-time analyses supported early diversification of calcar length in all cases, as evidenced by significantly low MDI values when compared to a null BM distribution (Fig. 2, Table 2). MDI values consistently increased when the calcar length disparity-through-time curve was compared to a distribution generated under an EB model of evolution.

**Figure 2.**
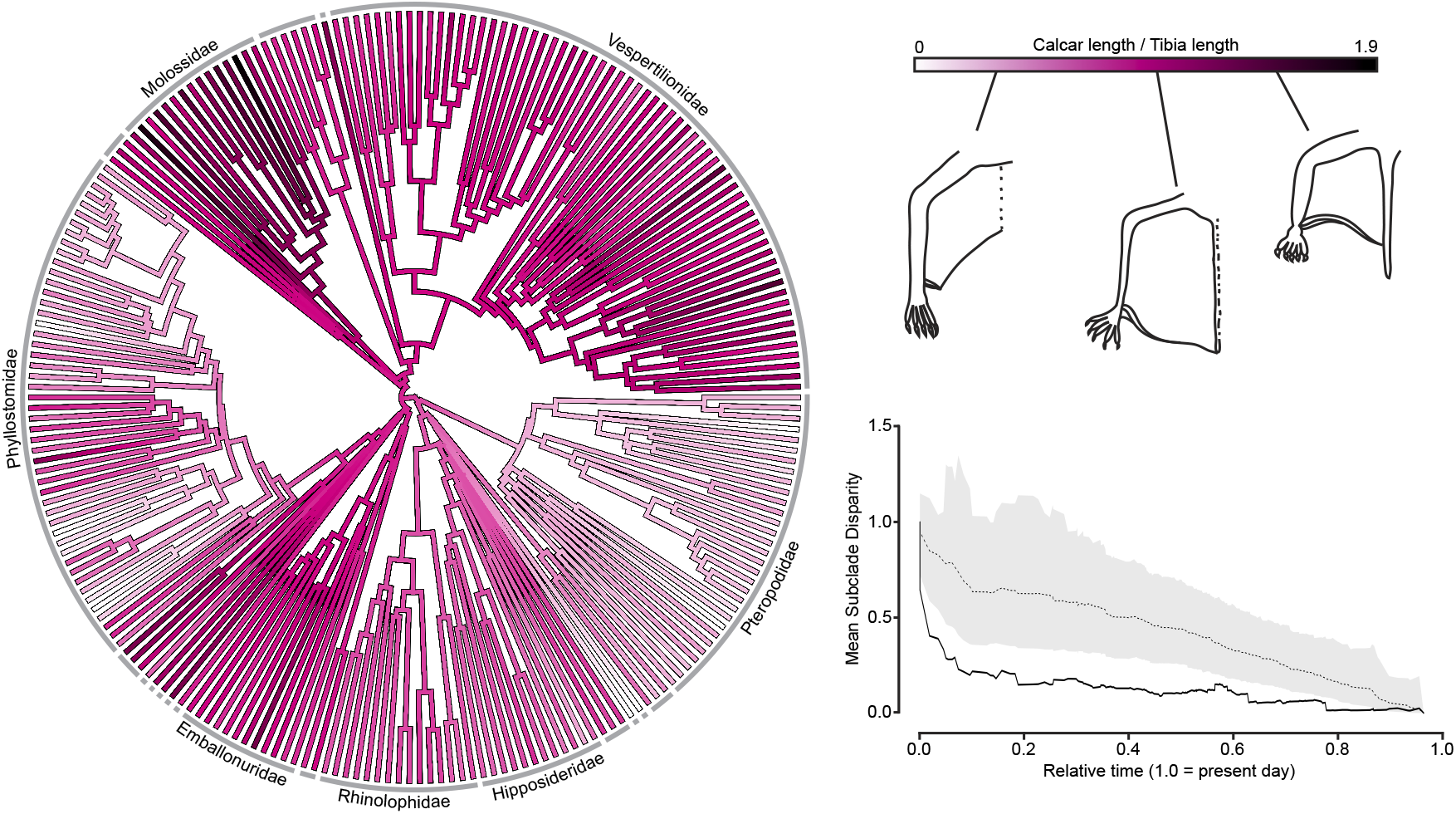
Relative calcar length varies extensively and diversified early in bat evolutionary history. Ratio of calcar length-to-tibia length is plotted on a phylogeny of Chiroptera. Gray lines around the phylogenetic tree designate bat families; species-rich families are labeled. Schematic drawing on the color scale illustrate representative hindlimb morphologies for different calcar lengths. In the diversity-through-time plot of mean subclade disparity, the black line indicates the mean subclade disparity through time for the measured calcar-to-tibia length ratios, the dotted line to 1,000 Brownian motion simulations, and the gray band a 95% confidence range for the simulations.

**Table 1.**
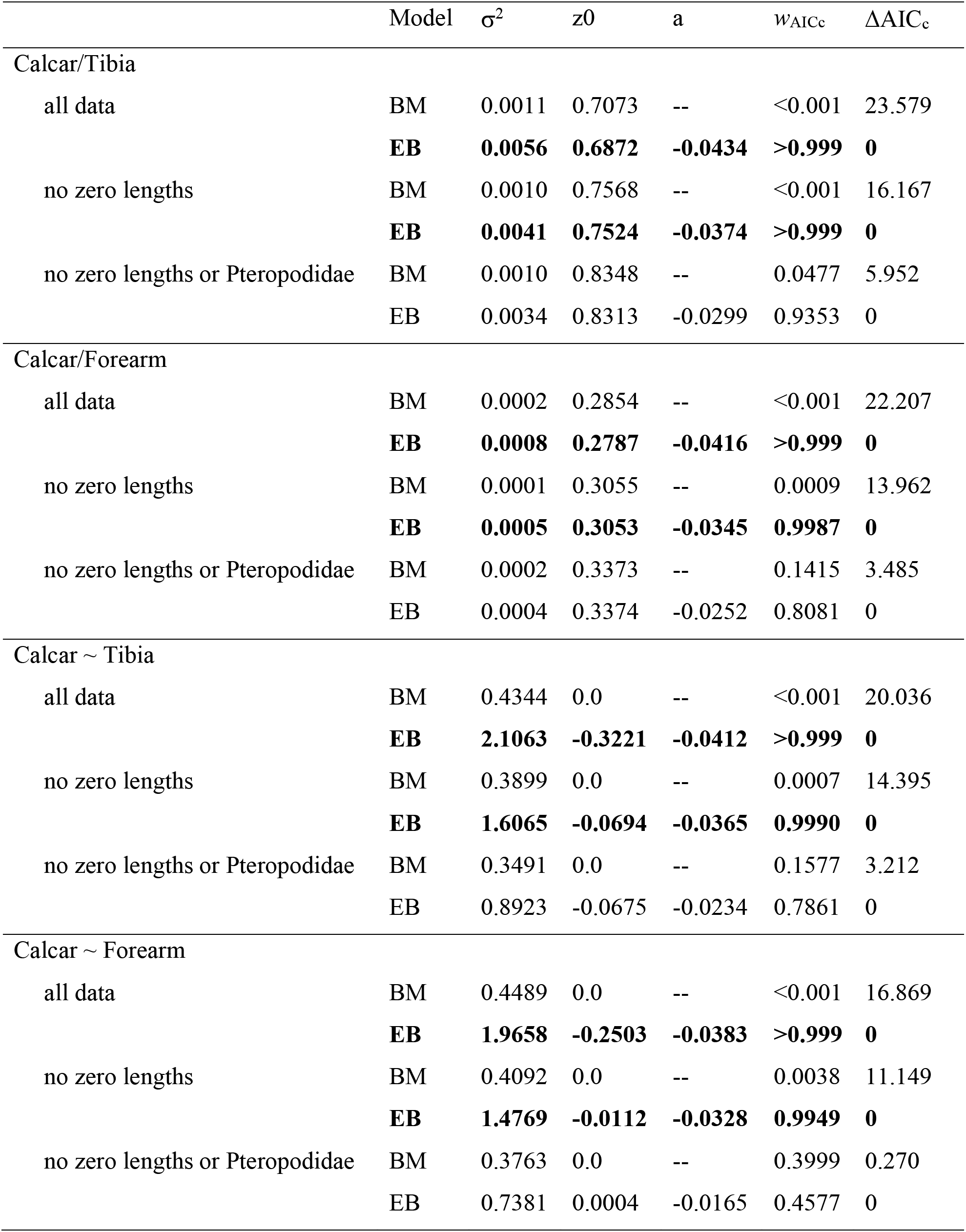
Results from evolutionary model comparisons. Calcar/Tibia and Calcar/Forearm indicate models considering ratios of calcar length to tibia and forearm length, respectively; Calcar~Tibia and Calcar~Forearm indicate models using residuals of phylogenetic regressions of the same variables. BM = Brownian motion model; EB = Early Burst model; a, σ^2^, and z0 are the fit parameters of those models corresponding to the names used in the “fitContinuous” function (see Material and Methods); *w*_AICc_ = AIC_c_ weights. All OU models collapsed to BM models, so only BM and EB results are shown. Bold text emphasizes models with *w*_AICc_ > 0.99.

| | Model | $\sigma^2$ | z0 | a | $W_{AICc}$ | $\Delta AIC_c$ |
| --- | --- | --- | --- | --- | --- | --- |
| **Calcar/Tibia** |  |  |  |  |  |  |
| all data | BM | 0.0011 | 0.7073 | -- | <0.001 | 23.579 |
|  | **EB** | **0.0056** | **0.6872** | **-0.0434** | **>0.999** | **0** |
| no zero lengths | BM | 0.0010 | 0.7568 | -- | <0.001 | 16.167 |
|  | **EB** | **0.0041** | **0.7524** | **-0.0374** | **>0.999** | **0** |
| no zero lengths or Pteropodidae | BM | 0.0010 | 0.8348 | -- | 0.0477 | 5.952 |
|  | EB | 0.0034 | 0.8313 | -0.0299 | 0.9353 | 0 |
| **Calcar/Forearm** |  |  |  |  |  |  |
| all data | BM | 0.0002 | 0.2854 | -- | <0.001 | 22.207 |
|  | **EB** | **0.0008** | **0.2787** | **-0.0416** | **>0.999** | **0** |
| no zero lengths | BM | 0.0001 | 0.3055 | -- | 0.0009 | 13.962 |
|  | **EB** | **0.0005** | **0.3053** | **-0.0345** | **0.9987** | **0** |
| no zero lengths or Pteropodidae | BM | 0.0002 | 0.3373 | -- | 0.1415 | 3.485 |
|  | EB | 0.0004 | 0.3374 | -0.0252 | 0.8081 | 0 |
| **Calcar ~ Tibia** |  |  |  |  |  |  |
| all data | BM | 0.4344 | 0.0 | -- | <0.001 | 20.036 |
|  | **EB** | **2.1063** | **-0.3221** | **-0.0412** | **>0.999** | **0** |
| no zero lengths | BM | 0.3899 | 0.0 | -- | 0.0007 | 14.395 |
|  | **EB** | **1.6065** | **-0.0694** | **-0.0365** | **0.9990** | **0** |
| no zero lengths or Pteropodidae | BM | 0.3491 | 0.0 | -- | 0.1577 | 3.212 |
|  | EB | 0.8923 | -0.0675 | -0.0234 | 0.7861 | 0 |
| **Calcar ~ Forearm** |  |  |  |  |  |  |
| all data | BM | 0.4489 | 0.0 | -- | <0.001 | 16.869 |
|  | **EB** | **1.9658** | **-0.2503** | **-0.0383** | **>0.999** | **0** |
| no zero lengths | BM | 0.4092 | 0.0 | -- | 0.0038 | 11.149 |
|  | **EB** | **1.4769** | **-0.0112** | **-0.0328** | **0.9949** | **0** |
| no zero lengths or Pteropodidae | BM | 0.3763 | 0.0 | -- | 0.3999 | 0.270 |
|  | EB | 0.7381 | 0.0004 | -0.0165 | 0.4577 | 0 |

**Table 2.** Results from disparity-through-time analyses. MDI = morphological disparity index; BM = Brownian motion model; EB = Early Burst model.

|  | MDI (BM) | MDI (EB) |
| --- | --- | --- |
| Calcar/Tibia |  |  |
| all data | -0.284 (p < 0.001) | -0.105 (p = 0.034) |
| no zero lengths | -0.275 (p < 0.001) | -0.119 (p = 0.036) |
| no zero lengths or Pteropodidae | -0.236 (p = 0.002) | -0.112 (p = 0.063) |
| Calcar/Forearm |  |  |
| all data | -0.287 (p < 0.001) | -0.113 (p = 0.023) |
| no zero lengths | -0.278 (p < 0.001) | -0.125 (p = 0.013) |
| no zero lengths or Pteropodidae | -0.223 (p = 0.001) | -0.113 (p = 0.065) |
| Calcar ~ Tibia |  |  |
| all data | -0.223 (p < 0.001) | -0.056 (p = 0.182) |
| no zero lengths | -0.221 (p = 0.001) | -0.066 (p = 0.161) |
| no zero lengths or Pteropodidae | -0.195 (p = 0.003) | -0.0898 (p = 0.108) |
| Calcar ~ Forearm |  |  |
| all data | -0.222 (p = 0.001) | -0.055 (p = 0.195) |
| no zero lengths | -0.207 (p = 0.004) | -0.0752 (p = 0.128) |
| no zero lengths or Pteropodidae | -0.17 (p = 0.027) | -0.0931 (p = 0.129) |

Detailed investigation of calcar anatomy with μCT scans revealed that bat ankles exhibit numerous tarsal modifications and collectively contain multiple accessory ossicles (Fig. 3; descriptions in Appendix S1). However, none of these osteological modifications refute the status of the calcar as a skeletal neomorphism or morphological novelty. In no bat species is the calcar contiguous with another tarsal, nor is the calcar an obviously repeated skeletal element. The calcar of any one bat species is only anatomically similar in both structure and location to calcars of other bats and not to another tarsal element.

**Figure 3.**
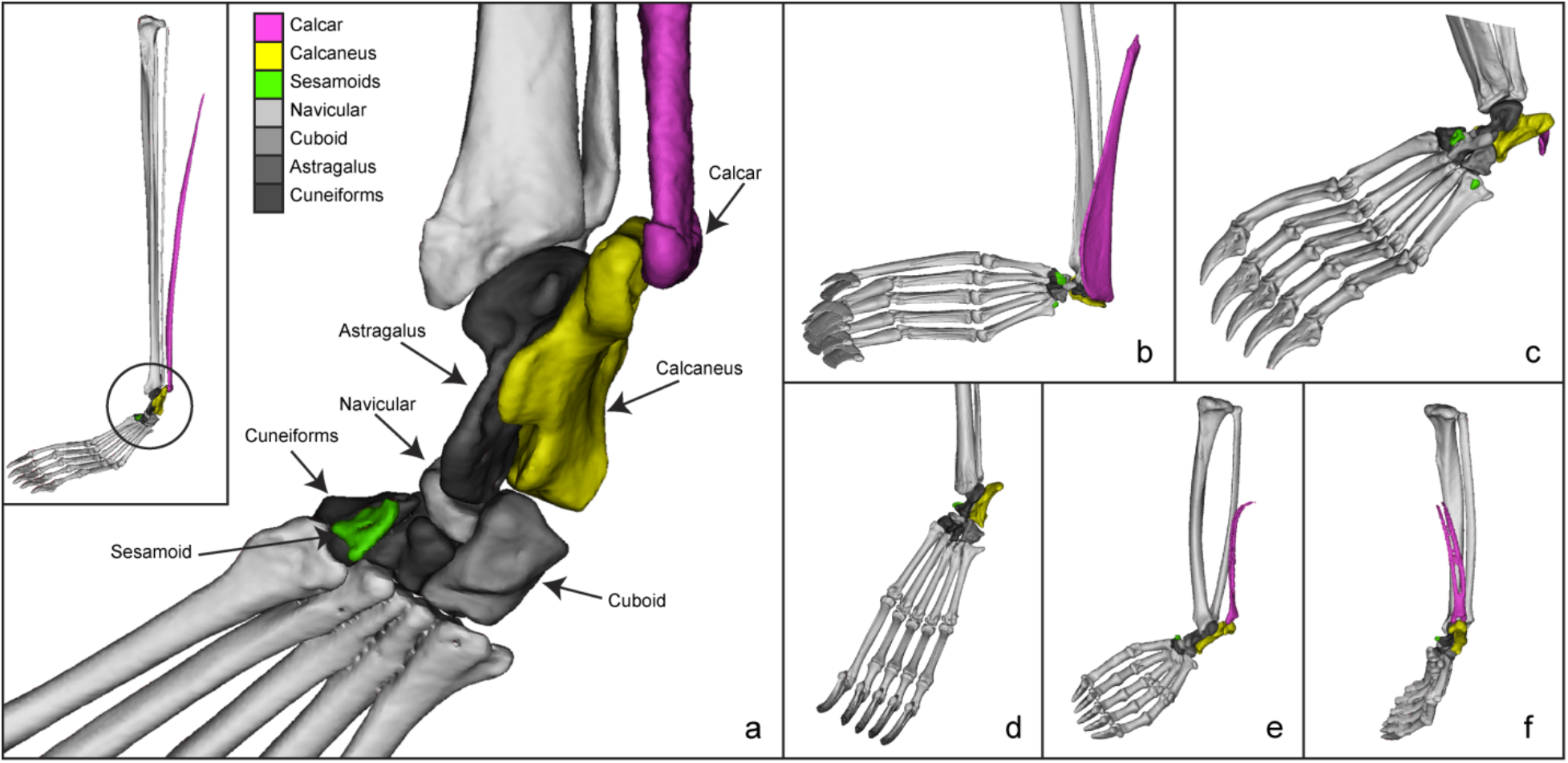
Bat ankle morphologies as demonstrated by rendered μCT scans. (a) Ankle of *Balantiopteryxplicata*, demonstrating calcar-calcaneus articulation (in pink-yellow), the other typical mammalian tarsals (in addition to the calcaneus; in shades of gray), and an additional sesamoid (in green). Inset demonstrates the ankle position relative to the full leg. Other bat feet μCT scans pictured are (b) *Noctilio leporinus*, (c) *Desmodus rotundus*, (d) *Rhinolophus affinis* (AMNH 234034; calcar not visible due to lack of calcification), (e and f) *Mystacina tuberculata* (MVZ 173918). All are pictured in plantar view except (f), which is medial to show calcified tines on calcar.

Histological sections complemented the μCT scans in revealing tissue-level diversity in bat calcars. While calcars are predominantly composed of uncalcified or calcified cartilage, some calcars contain ossified tissue (Fig. 4; Appendix S1). The calcar of *Noctilio leporinus* (Noctilionidae) is composed of thick cortical bone in the section proximal to the ankle, and both μCT scans and histological sections demonstrated the formation of trabeculae (Fig. 4a, b, c). The type of connective tissue also varies within a single calcar, along a continuum of cartilage, calcified cartilage, and bone. The calcar of *Molossus molossus* (Molossidae) is bony proximately and cartilaginous distally; as the bone grades into cartilage, only the interior of the calcar shaft is bony, and this bony tissue is surrounded by a thick layer of tissue that appears more cartilage-like (Fig. 4d, e). This partially bony calcar contrasts with the typical cartilaginous calcar of the other species, as exemplified by the primarily calcified cartilage calcar of *Eptesicus fuscus* (Fig. 4f). Both the *E. fuscus* and *M. molossus* calcars are surrounded by a thick, perichondrium-like envelope (Fig. 4e and 4f, respectively). *Pteronotus quadridens* (Mormoopidae) and *Macrotus waterhousii* (Phyllostomidae) also have bony proximal ends of their calcars, but the degree to which this ossification extends distally varies between the two species (Appendix S1). The short calcar of *Desmodus rotundus* (Phyllostomidae) also exhibits bony tissue (Fig. 4g).

**Figure 4.**
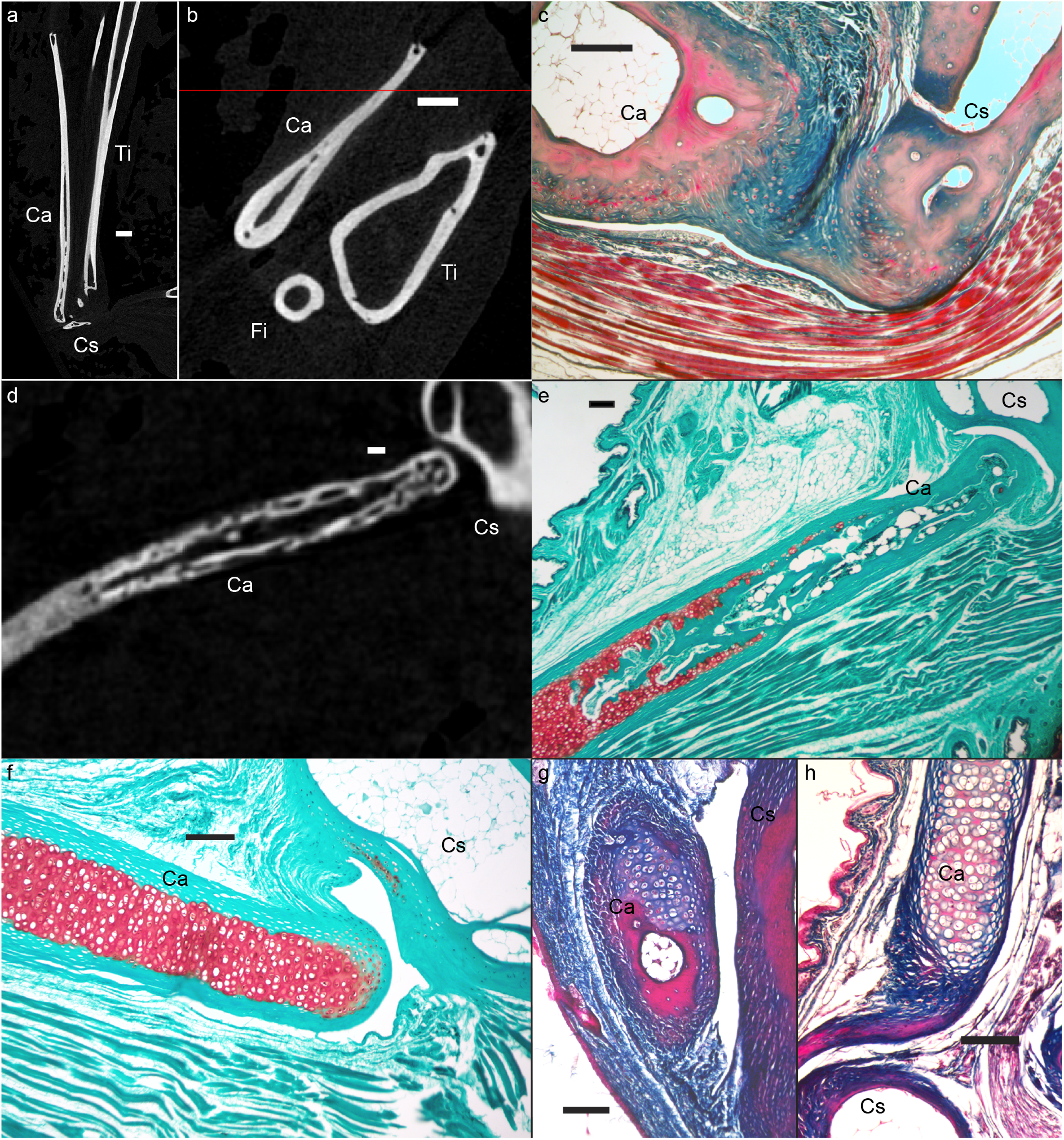
Histological diversity in the bat calcar. (a) Slice of μCT scan through the longitudinal axis of the calcar of *Noctilio leporinus.* (b) Axial μCT scan slice through the hindlimb of *N. leporinus*, demonstrating cross-sectional shapes of the calcar and leg bones. (c) Mallory-stained histological section through the ankle of *N. leporinus*, demonstrating bony calcar tissue and a ligamentous connection between the calcar and calcaneus. (d) Slice of μCT scan and (e) fast green/safranin O-stained histological section through the longitudinal axis of the calcar of *Molossus molossus*, demonstrating bony tissue in the calcar near the synovial joint with the calcaneus, which then transitions distally to calcified cartilage. (f) Fast green/safranin O-stained histological section of *Eptesicusfuscus*, showing a fully-cartilaginous calcar and a synovial joint between the calcaneus and the calcar. (g) Mallory-stained histological section of *Desmodus rotundus*, demonstrating bony nodule of calcar near the synovial articulation with the calcaneus. (h) Mallory-stained histological section demonstrating calcar presence in *Rhinopoma hardwickii* (FMNH 123185). Ca = calcar, Cs = calcaneus, Fi = fibula, Ti = tibia. In all sections the scale bar indicates 100μm, except for (b) and (b) where it is 500μm.

Histological sections also confirmed the presence of a synovial joint between the calcar and the calcaneus in several bat species (Fig. 4e, f, g; Appendix S1) and the presence of a relatively small, uncalcified, cartilaginous calcar in one species in which the calcar was thought to be absent (*Rhinopoma hardwickii*, Rhinopomatidae; Fig. 4h). Our anatomical analyses also highlighted known shape differences across bat calcars; although most calcars take the form of a rod with an approximately elliptical cross-section, some exhibit notably divergent shapes. For example, a cartilaginous hook-like “keel” structure protrudes from the main shaft of the calcar in some species, including *Eptesicus fuscus, Myotis californicus* (both Vespertilioonidae), and *Thyroptera tricolor* (Thyropteridae). The bony portion of the calcar of *Noctilio leporinus* exhibits an antero-posteriorly flattened cross-section with multiple cavities in the bony tissue (Fig. 4b). We describe, for the first time, that the calcar of *Mystacina tuberculata* (Mystacinidae) has two distinct calcified tines (Fig. 3e, f), a unique morphology among the calcars in our sample.

The diceCT scans and dissections of pteropodid feet revealed calcar anatomical diversity within the Pteropodidae. The diceCT scan of *Cynopterus brachyotis* indicates that the calcar and the tendon of the gastrocnemius muscle make two separate, distinct insertions on the calcaneal tuberosity. We confirmed this observation through a dissection in which we were able to cleanly pass a pin between the insertions of the calcar and the tendon on the calcaneus (Fig. 5). However, dissections of the calcars of *Rousettus aegyptiacus* and *Pteropus* sp. indicated that the calcar tissue is contiguous with the tendon of the gastrocnemius muscle. DiceCT scans of these species were inconclusive, as iodine solution only slightly increases CT scan image contrast in cartilage. More detailed anatomical descriptions of each species examined with μCT scanning and histological sectioning are provided in Appendix S1.

**Figure 5.**
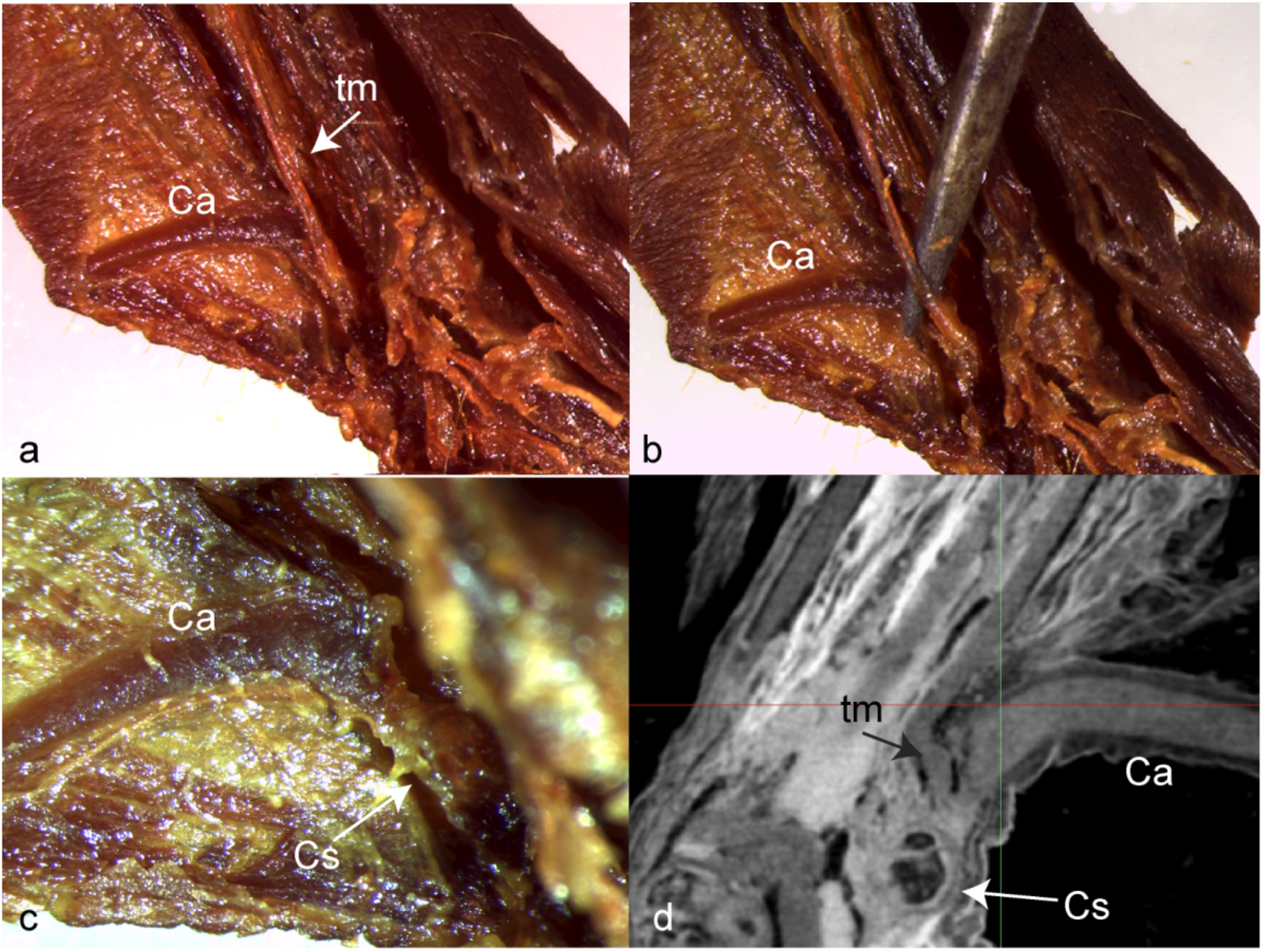
Photographs of dissection of the *Cynopterus brachyotis* ankle, demonstrating separation between the calcar and the tendon of the gastrocnemius muscle. (a) – (c) are dissection photos of an iodine-stained specimen. (b) pin demonstrates the separation between the calcar and the tendon. (c) shows the insertion of the calcar on the calcaneal tuberosity after the tendon has been dissected out. (d) is a slice of the diceCT scan demonstrating the separation between the calcar and the tendon and their two distinct insertions on the calcaneus. Ca = calcar, Cs = calcaneus, tm = tendon of the gastrocnemius muscle.

## DISCUSSION

The bat calcar is a skeletal novelty that has anatomically diversified widely throughout Chiroptera. This diversification appears to have occurred early in chiropteran history, as evidenced by support for an early burst model of calcar length evolution and the corresponding negative morphological disparity indices. This is consistent with evidence for early diversification of bats in the fossil record and an overall declining rate of speciation in Chiroptera (Shi and Rabosky 2015). Eocene bat postcrania are best preserved in the Green River Formation and the famous Messel Lagerstätten near Messel, Germany. Although *Onychonycteris finneyi* is known to have had a calcar, no calcars have been found in postcranial fossils of *Icaronycteris index*, a later Green River bat with limb proportions typical of some extant bats. Among the Messel bats, *Hassianycteris, Palaeochiropteryx*, and *Tachypteron* had calcars, but no calcars have been reported in specimens of *Archaeonycteris* (Simmons and Geisler 1998, Storch et al. 2002). Additionally, no evidence of an articulation facet has been found on the calcanei of *Icaronycteris* and *Archaeonycteris* (Simmons and Geisler 1998). Because calcars vary in amount of calcification, it is possible that uncalcified cartilage calcars were not preserved in these taxa; nonetheless, it is clear that Eocene bats exhibited diversity in either the presence of a calcar or in the amount of calcar calcification soon after the first bats evolved flight.

We found weaker support for the EB model when only non-pteropodid calcars were included in the analyses. However, our pteropodid diceCT scan and dissection results call into question the proposition that the pteropodid calcar is not homologous to the calcar of other bats. We have demonstrated that the calcar morphology of at least one pteropodid individual (*Cynopterus brachyotis*) differs from the calcar morphology of other pteropodids; its relation to the surrounding connective tissue makes it more similar to the “microbat” calcar condition. This intermediate anatomical condition in *C. brachyotis* suggests that it is more appropriate to consider the calcars of all bats in macroevolutionary analyses, rather than just those of the paraphyletic “microchiroptera.”

Support for the EB model of morphological evolution is notoriously low in the macroevolution literature (Harmon et al. 2010). It has been proposed that this could be an artefact of either hypothesis testing at too low of a taxonomic level, such that the signal of the “early burst” of the higher-level clade has been lost, or a consequence of testing variables that are not functionally-linked to the specific radiation, such as body mass and overall shape (Slater and Friscia, 2018). The evolution of wings in the early Chiroptera is a type of extensive morphological change that would be expected to precede a burst of diversification, as flight would allow access to an entirely new ecospace (other examples summarized in Erwin 2015). The calcar abruptly appeared in the fossil record as part of this wing structure and is now found in the vast majority of bats. When we tested an early burst hypothesis of calcar evolution across all of Chiroptera, we found that the calcar—a distinct synapomorphy associated with an aerial ecological mode—retains the signal of an early diversification burst. The true key innovation, however, is likely the full wing apparatus, which not only includes the novel calcar but also the elongation of the forelimb bones and the evolution of novel and developmentally-retained wing membranes. The functional relevance of the calcar within the wing is untested, although it is generally assumed that the calcar plays a role in supporting the hindlimb membrane during flight (Vaughan 1970).

Across extant bats, the calcar exhibits interspecific diversity in anatomical parameters that are likely to affect function, both in terms of overall structure (e.g., length and shape) and material (histological) composition. Although others have noted differences in the amount of calcar calcification among species based on dissection observations and clearing and staining procedures (Schutt and Simmons 1998, Koyabu and Son 2014, Reyes-Amaya et al. 2017), this is the first study to confirm the presence of ossified tissue in the bat calcar. Given that there is extensive variation in material properties between cartilage, calcified cartilage, and bone (Currey 2002), interspecific variation in calcar tissue composition, length, and/or shape would result in interspecific differences in responses to applied loading (e.g., muscular contraction or resistance of a stretched membrane). Bat hindlimbs play important functional roles in prey capture (Fish et al. 1991), roosting (Simmons and Quinn 1994), and possibly flight (Cheney et al. 2014). The anatomical variation described here suggests calcar function may vary across species with different ecologies that are subject to different functional evolutionary pressures. For instance, in *Myotis*, long calcars were found to be associated with a trawling foraging strategy (Fenton and Bogdanowicz 2002). Additionally, the presence of a synovial joint between the calcar and the calcaneus, in combination with the presence of skeletal muscles that insert on the calcar (Schutt and Simmons 1999, Glass and Gannon 1994, Stanchak and Santana 2018), suggests a kinematic functional role for the calcar. Although there are reported observations of moving calcars (e.g., in *Noctilio leporinus* as they trawl bodies of water for fish prey; Vaughan 1970, Altenbach 1989), calcar motion has not yet been confirmed with a rigorous kinematic analysis in any bat species. Further detailed, quantitative analyses of calcar biomechanics, including material testing and behavioral experiments, are required to estimate the magnitude of the effect of anatomical variation on any potential calcar function.

The origin of the calcar is still a mystery. It meets Hall’s (2015) definition of neomorphs, which *“seem to appear out of nowhere, de novo*, but are present in most if not all individuals of a species” as well as Müller and Wagner’s (1999) definition of a morphological novelty as “neither homologous to any structure in the species nor homonomous to any other structure of the same organism.” Although the immediate ancestry of the chiropteran lineage is unknown (Halliday et al. 2017), no calcar-like structure is found in earlier eutherian mammals. However, the discovery of a calcar in a Mesozoic mammaliaform (Meng et al. 2017) raises the possibility of a deep homological explanation for the origin of calcar (Shubin et al. 2009).

One proximate hypothesis for the origin of the calcar is that it initially develops within existing connective tissue in the hindlimb membrane via a process of metaplasia (Carter and Beaupré 2007). The condition of the pteropodid calcar, as described here, may provide incremental support for this hypothesis. Connective tissue (cartilage, tendon, and even bone) is both plastic and labile (Hall 2015). The calcar may have arisen in a mass of connective tissue in close proximity to the calcaneus, perhaps as that mass of tissue was placed under stress during the development of the hindlimb membrane. Consequently, differences among species in the association of the calcar with the calcaneus may be the result of relatively minor developmental alterations. Our finding of many sesamoids in bat feet, consistent with a recent assessment of bat sesamoids (Amador et al. 2018), suggests a propensity for metaplastic cartilage and bone development in bat feet, as tendon metaplasia is hypothesized to play a role in sesamoid development (Sarin et al. 2002; but see also Eyal et al. 2015, 2018). Developmental plasticity may also lead to intraspecific variation in calcar anatomy or even presence. This might be a fruitful path of further study in light of our finding of a small, calcar-like structure in the foot of one specimen of *Rhinopoma hardwickii.*

Anatomical structures of ambiguous homology are under-explored in studies of morphological evolution. The bat calcar is an anatomically diverse skeletal novelty found in a vast majority of species of a highly diverse clade of vertebrates. It evolved into a potentially functionally-important part of the bat wing, morphologically diversifying during the early radiation of bats. Additional, focused studies of the bat calcar—especially of its function and development—have a high potential to yield new knowledge of skeletal biology and a better understanding of the mechanisms through which the skeleton evolves into novel forms.

## Supporting information

Supplementary Information - Anatomical Descriptions

Supplementary Data - Used in Analyses

Supplementary Data - Raw

**Author Contributions:** KES and SES conceived of the project. KES collected and analyzed data. KES, JHA, and SES interpreted the data analysis and wrote the paper.

## Acknowledgements

**Acknowledgements:** We thank the Division of Mammals at the Smithsonian National Museum of Natural History, the Department of Mammalogy at the American Museum of Natural History, the Mammal Collection at the Museum of Vertebrate Zoology, the Slater Museum of Natural History, the Mammal Collection at the Field Museum of Natural History, the Herring Lab at the University of Washington, and the Lubee Bat Conservancy for providing access to specimens. L. Leiser-Miller collected many specimens in the Santana Lab collection. O. Okoloko helped with CT scans. L. Zeman, E. Johnson, and members of the Herring Lab advised on histological technique, and A.P. Summers provided use of the μCT scanner at Friday Harbor Laboratories. The project benefited from discussion with participants of the 2015 and 2018 Annual Meetings of the North American Society for Bat Research. A.A. Curtis, Z.A. Kaliszewska, R.M. Kelly, and L. Leiser-Miller provided advice on several manuscript drafts. KES was funded by a National Science Foundation (NSF) Doctoral Dissertation Improvement Grant (#1700845), a Theodore Roosevelt Memorial Grant from the American Museum of Natural History, a Grant-in-Aid of Research from the American Society of Mammalogists, the Iuvo Award from the University of Washington Department of Biology, and funds from the Department of Mammalogy at the Burke Museum of Natural History and Culture. SES and JHA were funded by NSF Grant #1557125.

**Data Accessibility Statement:** CT scans will be uploaded to Morphosource upon acceptance and available at time of publication. Data sets will be provided as Supporting Information associated with the publication.

