## Supplementary Information - Anatomical Descriptions for "Anatomical diversification of a skeletal novelty in bat feet"

This document includes:
Appendix S1 – Specimen list for μCT scanning and histology with CT scanner settings and anatomical descriptions.

Additional Supporting Information includes:

CalcarAnat_Evolution_SuppInfoRawData.csv (a list of specimens and measurements from which we compiled our calcar length data)

CalcarAnat_Evolution_SuppInfoAnalysisData.csv (data set used in calcar length macroevolutionary analyses)

Institution Acronyms in List of Specimens:
AMNH: American Museum of Natural History Mammal Collection
FMNH: Field Museum of Natural History Mammal Collection
Lab: Santana Lab in the Department of Biology at the University of Washington
MCZ: Museum of Comparative Zoology Mammal Collection
MVZ: Museum of Vertebrate Zoology Mammal Collection
NMNH: United States National Museum Mammal Collection at the Smithsonian National Museum of Natural History
PSM: Slater Museum of Natural History Mammal Collection

**Please see manuscript for full description of methods.**

**APPENDIX S1**

CT scanning and histology specimen list with anatomical descriptions of calcar and ankle modifications. Note that, because of the limited number of individuals per species in our sample, we were unable to confirm absence of a synovial joint in the histological sections. This is because the soft tissues around the joint did not always remain intact during sectioning, or the section was not in the adequate orientation/position to capture the tissues forming the synovial capsule.

*Artibeus jamaicensis* (Phyllostomidae) – Santana Lab 022614-25
Procedures: CT scanning and histological sectioning.
Small calcified nodule of cartilaginous calcar near articulation with calcaneus. No evidence of internal structure (i.e., not bony). Synovial joint between calcar and calcaneus. Sesamoid on plantar side of tibia, just above astragalus.
Scanned on Skyscan1172 at 8.88μm, 59kV, 167μA.

*Balantiopteryx plicata* (Emballonuridae) – Santana Lab 0229-06
Procedures: CT scanning and histological sectioning.
Long calcified portion of calcar with less densely-calcified or uncalcified interior. Calcar is entirely cartilaginous. More densely calcified "cap" to calcar at articulation with calcaneus. Sesamoid on plantar/lateral side of outer cuneiform (the cuneiform above MT1).
Scanned on Skyscan1172 at 7.97μm, 59kV, 167μA.

*Cynopterus brachyotis* (Pteropodidae) – Lubee Bat Conservancy
Procedures: CT scanning, diceCT scanning, dissection, and histological sectioning.
Uncalcified cartilage calcar not contiguous with the tendon of the *m. gastrocnemius*. Sesamoid on plantar/lateral side of outer cuneiform.
Scanned on Skyscan1172 at 10.58μm, 49kV, 200μA. diceCT scan on Skyscan1172 at 10.58μm, 40kV, 250μA after 2 days in 3% (total solute) Lugol's iodine.

*Desmodus rotundus* (Phyllostomidae) – Santana Lab 022714-06
Procedures: CT scanning and histological sectioning.
Calcar has small, bony nodule with clear internal structure but is distally cartilaginous. Synovial joint between calcar and calcaneus. Middle cuneiform is contiguous with/fused to the navicular. Sesamoid on plantar side of outer cuneiform and the middle cuneiform/navicular fusion. Sesamoid on plantar side of MT5.
Scanned on Skyscan1172 at 12.67μm, 59kV, 167μA.

*Eptesicus fuscus* (Vespertilionidae) – Santana Lab KES 037
Procedures: CT scanning and histological sectioning.
Cartilaginous calcar has lightly calcified section and an uncalcified "keel" hook. Synovial joint between calcar and calcaneus. Sesamoid on dorsal side of astragalus. Sesamoid on anterior/lateral side of outer cuneiform. Small nodule on plantar side of cuboid and under proximal end of calcar; these are possibly scan artefacts.
Scanned on Skyscan1172 at 11.50μm, 40kV, 250μA.

*Furipterus horrens* (Furipteridae) – AMNH 142903
Procedures: CT scanning (whole specimen).
Long, calcified cartilage calcar. Sesamoid above (superior to) outer cuneiform.
Scanned on Skyscan1173 at 12.14μm, 70kV, 114μA.

*Glossophaga longirostris* (Phyllostomidae) – Santana Lab GR110
Procedures: CT scanning and histological sectioning.
Uncalcified, cartilaginous calcar. Sesamoid on plantar side of tibia and astragalus.
Scanned on Skyscan1172 at 9.93μm, 59kV, 167μA.

*Hipposideros armiger* (Hipposideridae) – AMNH 272276
Procedures: CT scanning and histological sectioning.
Uncalcified cartilaginous calcar (but with perhaps a slight calcified band just above calcaneus). Sesamoid on dorsal side of astragalus. Sesamoid on plantar/lateral side of outer cuneiform. Sesamoid on plantar side of proximal end of $5^{th}$ metatarsal.
Scanned on Skyscan1172 at 9.80μm, 49kV, 200μA.

*Macrotus waterhousii* (Phyllostomidae) – Slater Museum FHA135
Procedures: CT scanning and histological sectioning.
Calcar is proximally bony. Sesamoid on plantar/lateral side of middle cuneiform. Lots of detritus in specimen that appears in scan.
Scanned on Skyscan1172 at 8.88μm, 59kV, 167μA.

*Megaderma spasma* (Megadermatidae) – FMNH 205388
Procedures: CT scanning and histological sectioning.
Uncalcified cartilaginous calcar (but with perhaps a slight calcified band just above calcaneus). Calcar forms synovial joint with calcaneus. Sesamoid on plantar/lateral side of outer cuneiform. Sesamoid on plantar side of proximal end of $5^{th}$ metatarsal.
Scanned on Skyscan1172 at 9.80μm, 49kV, 200μA.

*Molossus molossus* (Molossidae) – Slater Museum FHA1857
Procedures: CT scanning and histological sectioning.
Calcified calcar with internal bony structure at proximal end; cartilage distally. Calcar forms synovial joint with calcaneus. Sesamoid on dorsal side of astragalus. Sesamoid on plantar/lateral side of outer cuneiform. Sesamoid on plantar side of middle and outer cuneiforms.
Scanned on Skyscan1172 at 10.71μm, 59kV, 167μA.

*Myotis californicus* (Vespertilionidae) – Santana Lab KES 026
Procedures: CT scanning and histological sectioning.
Calcified cartilage calcar that forms an uncalcified “keel” hook. Calcar forms synovial joint with calcaneus. Sesamoid on plantar/lateral side of outer cuneiform.
Scanned on Skyscan1172 at 10.71μm, 40kV, 250μA.

*Mystacina tuberculata* (Mystacinidae) – MVZ 173918
Procedures: CT scanning (whole specimen).

Calcified calcar with internal structure that appears bony at proximal end. Calcification forms two long “tines”. Sesamoid on plantar/lateral side of outer cuneiform.

Scanned on Skyscan1173 at 20.96µm, 60kV, 133µA.

*Natalus mexicanus* (Natalidae) – Santana Lab 0229-02
Procedures: CT scanning and histological sectioning.

Calcified cartilage calcar with less densely-calcified or uncalcified interior. Medial and middle cuneiforms appear fused. Sesamoid on dorsal side of distal end of astragalus/proximal end of fused medial and middle cuneiforms.

Scanned on Skyscan1172 at 6.92µm, 59kV, 167µA.

*Noctilio leporinus* (Noctilionidae) – Slater Museum FHA1651
Procedures: CT scanning and histological sectioning.

Calcar has long calcified section that has distinctly bony internal structure, including what appear to be small trabeculae. Sesamoid on plantar side of outer and middle cuneiforms. Sesamoid on plantar side of proximal end of 5$^{th}$ metatarsal.

Scanned on Skyscan1172 at 12.15µm, 59kV, 167µA.

*Nycteris hispida* (Nycteridae) – USNM 478968
Procedures: CT scanning and histological sectioning.

Long, calcified cartilage calcar with less densely-calcified or uncalcified interior. Cuboid, medial cuneiform, and middle cuneiform are fused. Sesamoid on plantar side of outer cuneiform.

Scanned on Skyscan1172 at 6.88µm, 49kV, 200µA.

*Pteronotus quadridens* (Mormoopidae) – Slater Museum FHA 780
Procedures: CT scanning and histological sectioning.

Long, primarily calcified cartilage calcar with bony tissue at proximal end near articulation with calcaneus. Proximal cartilaginous portion of calcar has less-densely calcified interior. Sesamoid on plantar side of lateral outer cuneiform. Sesamoid on plantar side of proximal end of 5$^{th}$ metatarsal. Sesamoid on dorsal/medial side of cuboid.

Scanned on Skyscan1172 at 6.92µm, 59kV, 167µA.

*Pteropus sp.* (Pteropodidae) – Herring Lab #76.
Procedures: CT scanning, diceCT scanning, dissection.

Uncalcified cartilage calcar that is contiguous with the tendon of the *m. gastrocnemius* (based on dissection).

Scanned on Skyscan1172 at 26.44µm, 49kV, 200µA. diceCT scan on Skyscan1172 at 26.44µm, 40kV, 250µA after 3 days in 3% (total solute) Lugol’s iodine.

*Rhinolophus affinis* (Rhinolophidae) – AMNH 234034
Procedures: CT scanning and histological sectioning.

Uncalcified cartilaginous calcar. Sesamoid on dorsal side of distal end of astragalus.

Scanned on Skyscan1172 at 6.40μm, 49kV, 200μA.

*Rhinopoma hardwickii* (Rhinopomatidae) – FMNH 123185
Procedures: CT scanning and histological sectioning.

Uncalcified cartilaginous calcar. Sesamoid on dorsal side of distal end of astragalus. Sesamoid on plantar side of navicular. Sesamoid on plantar side of proximal end of 5$^{th}$ metatarsal.

Scanned on Skyscan1172 at 7.45μm, 49kV, 200μA.

*Rousettus aegyptiacus* (Pteropodidae) – Herring Lab #224.
Procedures: CT scanning, diceCT scanning, dissection, and histological sectioning.

Uncalcified cartilage calcar that is contiguous with the tendon of the *m. gastrocnemius* (based on dissection). Sesamoid on plantar/lateral side of outer cuneiform. Small nodule on dorsal side of foot between proximal ends of calcaneus and astragalus. This small nodule does not appear to be present in the scan of the other foot.

Scanned on Skyscan1172 at 13.22μm (Lft – analyzed in Mimics) 10.45μm (Rft), 49kV, 200μA.
diceCT scan (Rft) on Skyscan1172 at 10.18μm, 40kV, 250μA after 2 days in 3% (total solute) Lugol's iodine.

*Thyroptera tricolor* (Thyropteridae) – MVZ 158246
Procedures: CT scanning (whole specimen/hindlimbs only).

Long, primarily uncalcified cartilage calcar with calcified nodule at articulation with calcaneus. Calcar has two obvious uncalcified "keel"-like structures (based upon external examination of specimen). Navicular and middle cuneiform are fused. Sesamoid on plantar side of outer cuneiform.

Scanned on Skyscan1173 at 13.15μm (hindlimbs only in scan), 60kV, 133μA.
